# Cellular and clinical impact of Protein Phosphatase Enzyme epigenetic silencing in multiple cancer tissues

**DOI:** 10.1101/2022.03.03.482850

**Authors:** Ricky S. Joshi, Manel Esteller

## Abstract

Protein Phosphatase Enzymes (PPE) and protein kinases simultaneously control phosphorylation mechanisms that tightly regulate intracellular signaling pathways and stimulate cellular responses. In human malignancies, PPE and protein kinases are frequently mutated resulting in uncontrolled kinase activity and PPE suppression, leading to cell proliferation, migration and resistance to anti-cancer therapies. Cancer associated DNA hypermethylation at PPE promoters gives rise to transcriptional silencing (epimutations) and is a hallmark of cancer. Despite recent advances in sequencing technologies, data availability and computational capabilities, only a fraction of PPE have been reported as transcriptionally inactive as a consequence of epimutations. Using the Infinium HumanMethylation450 BeadChip, we compared DNA methylation profiles from 705 cancer patients across 5 major tissues and 3 cancer cell models against a cohort of healthy controls. Here, we report epimutations in PPE (and their interacting proteins or PPEIP) are a frequent occurrence in the cancer genome and manifest independent of transcriptional activity. We observed that different tumors have varying susceptibility to epimutations and identify specific cellular signalling networks that are primarily affected by epimutations. Additionally, RNA-seq analysis showed the negative impact of epimutations on most (not all) Protein Tyrosine Phosphatase transcription. Finally, we detected novel clinical biomarkers that inform on patient mortality and anti-cancer treatment sensitivity. We propose that DNA hypermethylation marks at PPEIP frequently contribute to the pathogenesis of malignancies and within the precision medicine space, hold promise as biomarkers to inform on clinical features such as patient survival and therapeutic response.

## Introduction

Protein phosphorylation is a post-translational modification that is vital for controlling signaling pathways and homeostatic maintenance of human cells such as metabolism, transcription/translation and cell division (Cohen, 1989). Protein phosphatases (PPE) are enzymes that catalyze the removal of phosphate groups from cell signaling proteins by hydrolysis, reversing the action of protein kinases (PK). The activities of both are highly coordinated and orchestrate responses from external stimuli and/or relay information for key transcriptional responses that stimulate or inhibit growth and protect the cell which are highly sensitive to small changes in PPE activity (Stebbing et al., 2014; Tonks, 2006). PPE are categorized into two subtypes according to their specificity and action; Protein Tyrosine Phosphatases (PTP) and Protein Serine/Threonine Phosphatases (PSTP). PTP are further divided into (a) receptor-like and non-receptor PTP and (b) Dual-Specificity Phosphatases (DUSP). Typical PTP catalyze phosphotyrosine residues whereas DUSP dephosphorylate phosphoserine/threonine and phosphotyrosine amino acids. PSTP are also divided into two sub-groups; PhosphoProtein Phosphatases (PPP) and Metal-dependent Protein Phosphatases (PPM). (Barford et al., 1998; Li et al., 2013). PPE and PK represent arguably the most studied group of proteins in the literature to date. The uncontrolled suppression of PPE has often been described as a cause of dysregulated cell signaling programs that lead to human disease (Bononi et al., 2011; Tonks, 2006). In cancer, an imbalance in phosphorylation equilibrium has been reported as a cause of abnormal cell proliferation, dissemination and insensitivity to therapeutic treatment (extensively reviewed in Turdo et al., 2021). Many PK are well-known oncogenes, therefore PPE that counteract PK are assumed to be tumor suppressors (Fontanillo and Köhn, 2016). To date, numerous PPE have been reported with tumor suppressor activity, PTEN as the most documented example. Identified as a tumor suppressor in 1997, *PTEN* was initially observed to be deleted in brain, breast and prostate cancer tissue (Li et al., 1997). Protein Phosphatase 2A (PP2A) is the most expressed PSTP and described as a tumor suppressor in several cancers, e.g. breast, lung and melanomas (Calin et al., 2000) due to its role in inhibiting signal transduction pathways such RAF-MEK-ERK and Ras/PI3K/PTEN/Akt/mTOR. These pathways favor a number of cellular functions vital for tumor growth, such as cancer proliferation, reduced sensitivity to apoptotic signals and activation of pro-survival pathways. (Steelman et al., 2011; Zhang and Claret, 2012). This dysregulation is essential to create an environment conducive for malignancies. PTP are one of the most recognized group of genes that make up the tumor suppressor family and are frequently inactivated and/or mutated in a variety of cancers (FUNATO et al., 2011; Wang, 2004). For example, PTPRT and PTPRD negatively regulate the JAK/STAT pathway (Veeriah et al., 2009). PTPRH and PTPRB suppress downstream signaling of PI3K/Akt/mTOR and MEK/MAPK pathways by dephosphorylation of the epidermal growth factor receptor (EGFR) (Yao et al., 2017) and PTPN13 in non-small cell lung cancer by the phosphorylation control of EGFR and HER2 (Scrima et al., 2012). As well as the PTP, several DUSP are crucial regulators of MAPK proteins, such as ERK and JNK and therefore considered critical tumor suppressors (Caunt and Keyse, 2013). Other types of phosphatases not discussed above also have important regulatory roles as tumor suppressors such as cellular prostatic acid phosphatase (PAcP) whose loss of expression leads to prostate carcinogenesis (Muniyan et al., 2014) or INPP5K, a phosphoinositide phosphatase gene associated with tumor suppressor activity in endometrial carcinoma (Hedberg Oldfors et al., 2015).

Epigenetic dysregulation has been identified in cancer cells and mainly consists of global DNA hypomethylation with DNA hypermethylation at promoters of specific tumor-suppressor genes resulting transcriptional silencing (Esteller, 2008). The clinical implications of DNA methylation has been an important characteristic to understand cellular transformation and is an important tool for cancer diagnosis, prognosis and therapy monitoring (Ortiz-Barahona et al., 2020). In spite of the overwhelming evidence for PPE as tumor suppressors and the role of DNA hypermethylation plays in transcription silencing, only a handful of PPE have been epigenetically characterized in primary tumors (Barazeghi et al., 2019; Li et al., 2017; Luo et al., 2016; Schmid et al., 2015; Takane et al., 2014; Tögel et al., 2018; Vanaja et al., 2006; Ying et al., 2006).

Advances in sequencing technologies, data availability and bioinformatic approaches have enabled simultaneous analyses of large sample sizes and -omic datasets. In this study, we performed an exhaustive, systematic analysis of promoter DNA methylation of genes that encode PPE and PPE-interacting proteins (PPEIP) in five different malignant tissues to identify aberrant DNA hypermethylation profiles that are absent in healthy controls (epimutations). We report the frequency of epimutations discerned in Colorectal, Esophageal, Lung, Pancreatic and Stomach cancers and their effect on transcription, gene regulatory network/pathways as well as providing examples of their clinical implications such as survival and response to treatment.

## Results

### Pan-cancer promoter hyper-methylation analysis of Protein Phosphatase Enzymes and Interacting Proteins

To identify epigenetic defects in Protein Phosphatase Enzymes and Interacting Proteins (PPEIP), we studied the DNA methylation status at promoter regions of 523 PPEIP genes (Supplementary data S1) in five cancer subtypes (Colorectal, Esophageal, Lung, Pancreatic and Stomach cancers) across three cell models; primary tumors (The Cancer Genome Atlas (TCGA), cancer cell lines (Iorio et al., 2016) and 3D embedded cancer cell cultures or organoids (Joshi et al., 2020). DNA Methylation profiling for 705 cancer samples was performed using the Illumina Infinium Human Methylation 450 BeadChip (450K) and compared against a baseline control cohort comprising 42 unrelated healthy individuals from the same five tissue types mentioned above (epimutations). A list of all datasets, sample IDs and primary tissues can be found in Supplementary data S2 (also see methods). A simplified overview of the analysis workflow is provided in Figure 1. The 450K platform allows for genome-wide CpG methylation analysis at 485,000 CpG’s located at various genomic regions including 98% of all promoters in refseq-annotated genes. In order to dispel potential probe representations issues, we first analyzed the distribution of probes at PPEIP promoters against all other 450K represented genes. 6296 450K probes informed CpG DNA methylation levels in PPEIP promoters (12 probes per gene) and 3260 (52%) of those, in CpG islands associated with PPEIP promoters (6 probes per CpG island) (Figure 2A). In comparison, 168,664 probes were annotated to all gene promoters (9 probes per gene) and 79,008 (47%) to all promoter-associated CpG islands (4 probes per gene) indicating a higher representation of probes for PPEIP gene promoters compared to all other gene promoters (Figure 2A). The definition of “promoter probes” were those probes that provided semi-quantitative methylation values for CpG’s located within 1500bp and 200bp of the transcription start site (TSS1500, TSS200), 5’ untranslated region (5’UTR) or 1^st^ exon of each gene. The majority of PPEIP gene promoter probes were found in the TSS1500 (30%) and the least in the 1^st^ exon (13%) (Figure 2B). Half of all PPEIP probes were distributed between the 5’UTR (29%) and the TSS200 (21%) (Figure 2B). The median number of probes per gene was 11, and 7 when considering only CpG islands in PPEIP promoters (Figure 2C). The Protein Tyrosine Phosphatase gene *PTPMT1* was the PPEIP with the most probes associated with its promoter (*n* = 60).

**Figure 1:**
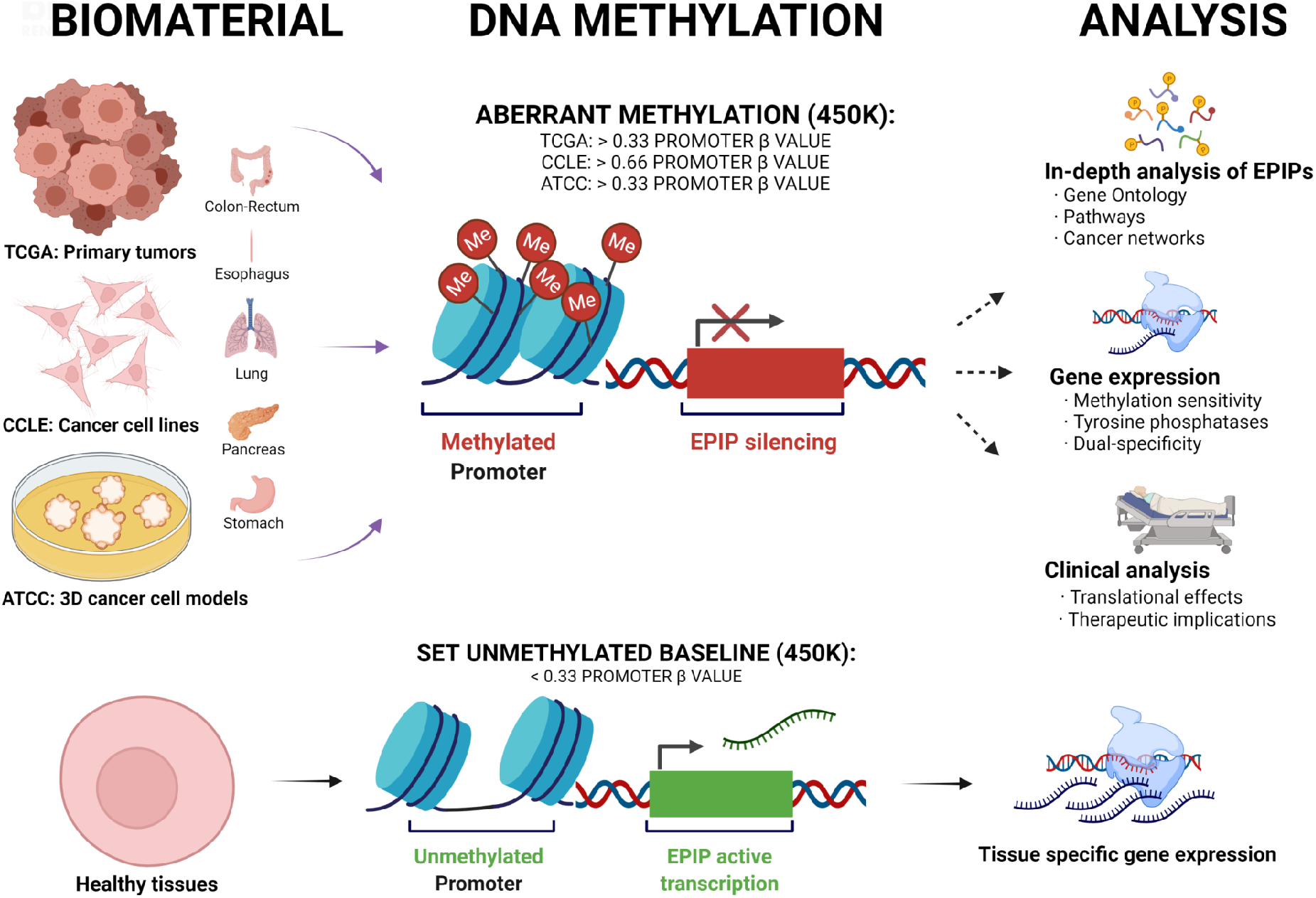
Analysis workflow for DNA methylation interrogation of Protein Phosphatase Enzymes and interacting protein (PPEIP) gene promoters. As a hallmark of cancer, cancer-specific increases in gene promoter DNA methylation result in transcription repression of key cancer-related genes. Here, publicly available genome-wide DNA methylation datasets (450K) from 5 cancer subtypes (Colorectal, Esophageal, Lung, Pancreatic and stomach cancers) were examined to discern aberrant methylation profiles that disrupt PPEIP expression and their related cellular pathways. Cancer samples were analyzed from primary tumors, cancer cell lines and 3D cancer cell models. Average promoter methylation beta values of >0.33 (primary tumors and organoids) and 0.66 (cancer cell lines) were identified and their effect on transcription, cell biology and translation were reported. DNA methylation profiles from concordant healthy samples were used as a baseline to determine cancer-specific DNA methylation-induced PPEIP silencing.

**Figure 2:**
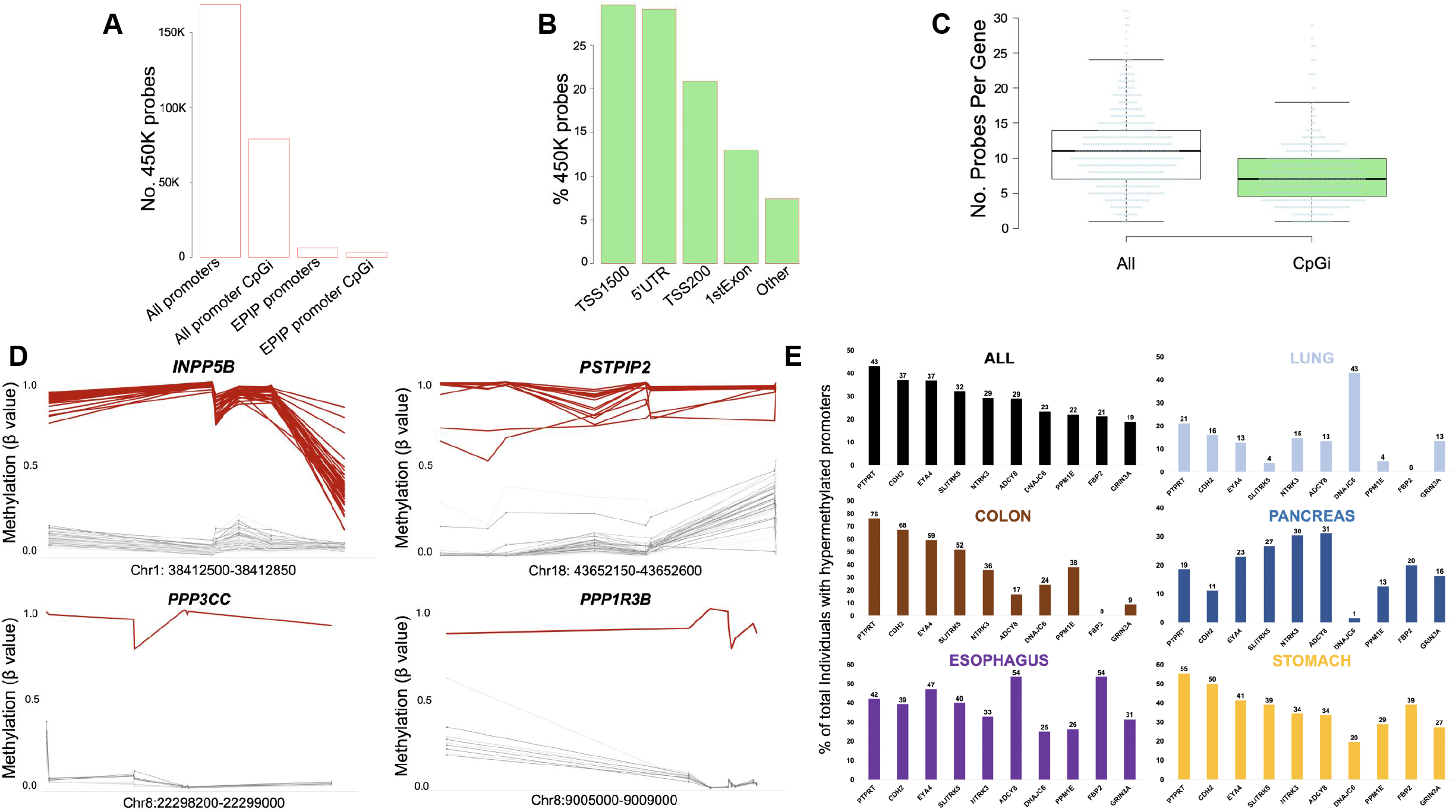
Infinium Human Methylation 450K BeadChip (450K) probe distribution and promoter hyper-methylation analysis reveals cancer-associated epimutations occur with varying frequencies in multiple tissues. A. Distribution of 450K probes across all gene promoters (and CpG island-specific) and PPEIP promoters (also showing CpG island specific probes). B. % of promoter CpG genomic annotations. TSS1500 = CpG present within 1500 bp from transcription start site (TSS), TSS200 = within 200 bp of TSS, 5’UTR = CpG is found within the 5’ untranslated region of the PPEIP and 1stExon = CpG found in the first exon of the PPEIP gene. The annotation “others” refers to promoter CpG’s that annotate to other genomic regions (such as gene bodies or introns) of overlapping genes. C. Distribution of promoter probes per gene annotated to both CpG islands and non-CpG islands. D. Examples of hyper-methylated PPEIP promoters for both recurrent (observed in >5%of the cancer population) and rare (<1% % of the cancer population) epimutations. Red lines present individuals with outlier epimutations and grey lines, healthy controls. E. Bar graph showing the ten most epimutated PPEIP genes. Numbers on the y-axis (and above each bar) represent the % of total cancer individuals in all cancer tissues (ALL) as well as a breakdown of epimutations identified in individuals of specific cancer related tissue.

### PPEIP promoter hyper-methylation profiles reveal distinct tissue susceptibility and frequency associated with cancer

Overall, 5007 hypermethylated Protein Phosphatase Enzymes and Interacting Proteins (PPEIP) gene promoters (epimutations) were identified in 593 cancer samples (84%), a median of 4 cancer-associated epimutations per individual. The distribution of epimutations across samples was significantly imbalanced (*P* = 2.2 10^-16^). Stomach cancer patients demonstrated the highest percentage of PPEIP epimutation distribution of 27% however, stomach cancer case only made up 18% of the total sample number (Supplementary Figure 1A). This represented the highest ratio of epimutation vs sample share, or observed epimutation distribution ratio (OEDR) of 0.59 (expected OEDR = 0) (Supplementary Figure 1B). Lung cancer patients showed with the lowest epimutation share with 9%, although 22% of all samples were from lung malignancies (−1.25 OEDR) (Supplementary Figure 1A and B). Colorectal, esophageal and stomach cancers showed positive OEDR (0.59 – 0.25) while lung and pancreatic cancer patients showed negative OEDR (−1.25 and −0.68 respectively) (Supplementary Figure 1B). 593 cancer patients (84%) presented with at least one hyper-methylated PPEIP promoter in 160 (31%) PPEIP genes analyzed as compared with healthy controls. Esophagus represented the cancer tissue with the highest number of individuals with at least one epimutation (98%) (Supplementary Figure 2A). This slight inconstancy with the OEDR stomach cancer data can be explained by the individual epimutation frequency in both tissues. 44% of all epimutations detected in the top 10% of individuals with the most epimutations accumulated were attributed to stomach cancer patients compared to only 28% in the esophagus (Supplementary data S3). This suggests that a subset of stomach cancer patients accumulate higher quantities of epimutations in fewer individuals as compared with esophageal cancer patients where epimutations are procured less inter-individually but more consistently across individuals. Lung cancer patients presented with the least number of epimutations (67%) consistent with OEDR data (Supplementary Figure 2A). 96% of all organoid samples harbored at least one hyper-methylated PPEIP promoter compared to 86% and 77% of all cancer cell lines and primary tumors respectively (Supplementary Figure 2B). To assess the data further, we applied our epimutation detection pipeline to 450K data from a separate test cohort of 47 control individuals presenting the five analyzed tissues to identify epimutations in a population of healthy individuals. 11 individuals (23%) were identified carrying 79 hyper-methylated PPEIP promoters as compared to 84% in cancer samples. A median epimutation count of 0 per individual was observed (Supplementary Figure S3A) as compared to 4 in the same tissues in a cancer context (Supplementary Figure S3B). Epimutations we detected were overwhelmingly enriched in cancer patients as compared to healthy individuals (P = 7.02 ×10^-18^). Interestingly, epimutations detected in healthy samples were only observed in two tissues (Esophagus *n*=7 and Pancreas *n*=4 individuals) (Supplementary data S4). This gave us confidence that the epimutations we identified through our bespoke bioinformatic pipeline were cancer-associated.

Next, we examined the frequency of PPEIP-associated epimutations in cancer patients. 5007 epimutations were detected in 160 PPEIP genes (Supplementary data S5). Of the 160 PPEIP genes, 88 (55%) were considered “rare” (observed in <1% of all cancer cases) and 41 (26%) as “recurrent” (identified in >5% of cancer samples) (Supplementary data S5). Many recurrent genes were known tumor suppressors with previously described cancer-related promoter hypermethylation anomalies (*PTPN13*, *DUSP5*, (Stebbing et al., 2014) *PPP1R14A* (Li et al., 2017), *PPP1R3C* (Takane et al., 2014), *PTPRM* (Barazeghi et al., 2019) and *IGFBP3* (Kawasaki et al., 2007; Torng et al., 2009; Ye et al., 2016) validating the robustness of our approach. We also detected several PPEIP genes with previously undescribed recurrent methylation changes (Figure. 2D). INPP5B (Inositol Polyphosphate-5-Phosphatase) is an anti-apoptotic protein with a proliferative role in different cancer types (H et al., 2013; J et al., 2009) and epimutations were observed in 130 individuals in all 5 tissues (Figure 2D). Proline-serine-threonine phosphatase interacting protein 2 or *PSTPIP2* promoter hyper-methylation was observed in 37 cases in all 5 tissues examined (Figure. 2D). PSTPIP2 is required for correct cell cycle function and dysregulation of PSTPIP2 contributes to abnormal proliferation and terminal differentiation in megakaryocytes (L et al., 2014). In addition to recurrent differentially methylated promoters, several patients displayed rare, previously undescribed epimutations at PPEIP promoters. For example, an epimutation was detected at the promoter of *PPP3CC* (Protein Phosphatase 3 Catalytic Subunit Gamma) in 3 individuals, colorectal (*n*=1) and stomach (*n*=2) cancer. PPP3CC repression contributes to invasion and growth of glioma cells (H et al., 2018). As loss-of-function genetic mutations in *PPP1R3B* gene have been associated with lung cancer (Kohno et al., 1999), similarly, DNA methylation associated transcription silencing mimic loss-of-function properties. We observed one individual with esophageal cancer phenotype harboring an epimutation at the *PPP1R3B* promoter (Figure. 2D).

We examined highly epimutated PPEIP genes and their described roles in human malignancies. Collectively, *PTPRT*, *CDH2*, *EYA4*, *SLITRK5*, *NTRK3*, *ADCY8*, *DNAJC6*, *PPM1E*, *FBP2* and *GRIN3A* were identified as those genes where most cancer individuals were observed to harbor hyper-methylated PPEIP promoters. An even distribution of epimutation count was not observed across all cancer tissue types consistent with our OEDR data. A breakdown of this is presented in Figure 2E. *PTPRT* was the most epimutated PPEIP with 43% of all cancer cases showing epimutations in this gene. PTPRT is a tyrosine phosphatase with a previously described role as a tumor suppressor in colorectal cancer (Wang, 2004). Interestingly, the authors also demonstrated that the most frequent genetically mutated tyrosine phosphatase gene was *PTPRT* in their colorectal cancer (CRC) cohort. In line with this finding, *PTPRT* was also the most epimutated gene in our CRC cohort; 76% of all CRC patients carried an epimutation. On the contrary, only 21% and 19% of all lung and pancreatic cancer patients respectively carried an epimutation in *PTPRT* (Figure 2E). Several of the highest epimutable genes have disparate prevalence of hypermethylated promoters between cancer tissue types. Eyes absent 4 (EYA4) is a threonine-tyrosine phosphatase (Okabe et al., 2009) previously described as a tumor suppressor in multiple cancers examined in this study (CRC (Kim et al., 2015), esophagus (Luo et al., 2018), lung (Wilson et al., 2014) and pancreas (Mo et al., 2016)). EYA4 promoter DNA methylation has been reported to be negatively correlated with gene expression and plays an important role in cell proliferation inhibition via Wnt and MAPK signaling pathways (Miller et al., 2010). The frequency of epimutation is highly contrasting as 59% of CRC patients carry hypermethylated EYA4 promoters compared with only 13% of lung cancer patients (Figure 2E). Another example is the NTRK3 gene. NTRK3 has been described in important cancer related pathways that promote both survival and cell proliferation and so its role as an oncogene (Vaishnavi et al., 2015) and tumor suppressor (Genevois et al., 2013) is not unexpected. We observed at least a twofold difference in individuals with NTRK3 hyper-methylated promoters between lung cancer (15%) and the other 4 cancers (CRC; 36%, stomach; 34%, Esophagus; 33% and Pancreas 30%) (Figure 2E).

### Pan-cancer promoter hyper-methylation of PPEIP affect cellular pathways and networks that favor tumor success

To further decipher the role of the 160 cancer-associated promoter hypermethylation susceptible PPEIP genes, we performed an enrichment analysis for gene networks, cellular pathways and transcription factor (TF) binding (Figure 3). For gene network and cellular pathway analysis, three highly cited software were used; Biocarta (Nishimura, 2001), Kyoto Encyclopedia of Gene and Genomes (KEGG) (Kanehisa et al., 2016) and curated WikiPathways (Pico et al., 2008). All three software demonstrated high overlap of well-known pathways described in cancer cells such PI3K-AKT (Vara et al., 2004), MAPK (Dhillon et al., 2007) and cellular metabolism (Seth Nanda et al., 2020) (Figure 3A,C,D). All three present actionable targets for anti-cancer drugs (Hennessy et al., 2005; Lee et al., 2020; Ngoi et al., 2020). Other interesting pathways include angiogenesis related VEGFA-VEGFR2 signaling (Zhao et al., 2015), regulatory circuits of STAT3 signaling pathways (Yu et al., 2014) as well as gene networks involved in aging (Hoeijmakers, 2009). We also interrogated transcription factor (TF) targets computed from ChIP-seq data from the ENCODE project (Sloan et al., 2016). The genes most affected by promoter methylation are also targets for TF that are highly mutated in cancer such as chromatin remodelers (EP300, HDAC2, KDM4A among others) (extensively reviewed in (Nair and Kumar, 2012)), cell cycle regulator (SIN3A (Shi and Garry, 2012)) and cell proliferation (YY1 (Gordon et al., 2006)) (Figure 3B).

**Figure 3:**
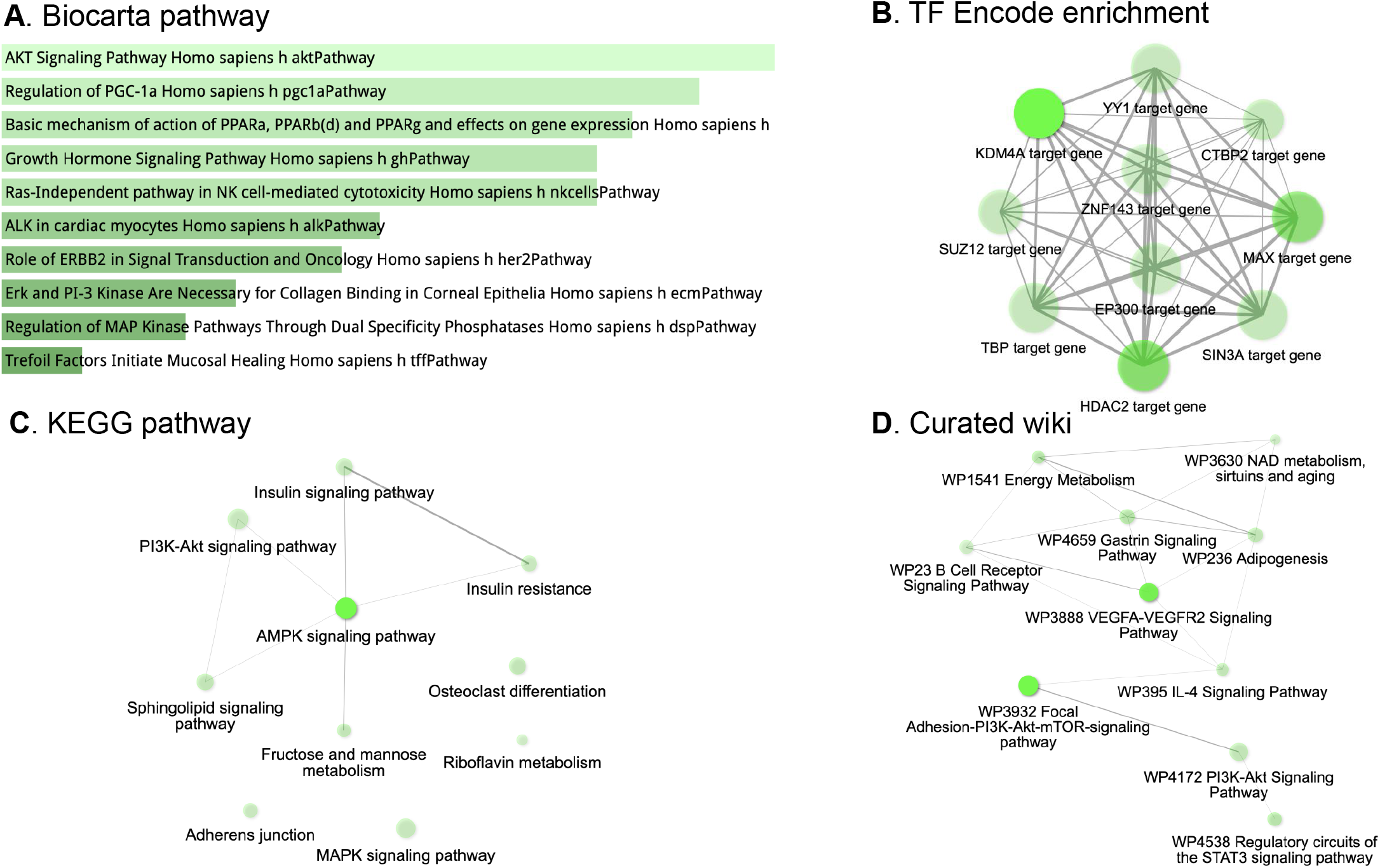
PPEIP genes with hyper-methylated promoters are involved in several cancer-associated cellular pathways and mechanisms. A. Biocarta pathway analysis for pathway gene enrichment. The shade of green presents the significance of the specific gene-set or term. B. Enrichment of transcription factor (TF) binding associated with hyper-methylated PPEIP genes from ChIP-seq experiments of >300 TFs from the ENCODE project. Bright green presents higher number of genes. C-D. Orthogonal cellular pathway analyses from two independent resources; Kyoto Encyclopedia of Genes and Genomes (KEGG) and Curated wiki. For Figures A-D, the brighter the tone of green, the more significant the term is. In all network images (B-D), the grey lines represent gene content similarity.

### Pan-cancer examination of epimutations in protein tyrosine and dual specific phosphatases exhibit aberrations in transcriptomic profiles affecting key cellular networks related to cancer

A number of Protein Tyrosine Phosphatases (PTPs) have been described in human cancers as tumor suppressor genes (Stebbing et al., 2014) and among the most studied include *PTPRM;* (Sudhir et al., 2015; Sun et al., 2012), *PTPN13;* (Scrima et al., 2012) and PTPRG; (Shu et al., 2010). Therefore, in primary tumors, PTPs would represent a subset of key genes where the effects of promoter DNA methylation induced transcriptional silencing are detrimental. In this regard, we closely examined PTPs for methylation sensitivity in primary tumors, cancer cell lines and 3D embedded cell cultures (organoids). We also performed the same analysis in dual-specificity phosphatases (DUSP) given their activity and role in cancer (Gao et al., 2021). Only few individuals presented hyper-methylated promoters for serine / threonine phosphatases (<2%) and therefore we focused our attention on PTP and DUSP genes. A list of PTPs and DUSPs was compiled from Ensembl and DNA methylation profiles generated for 43 PTP and 24 DUSP genes (Supplementary data S6). 17 PTP (40%) and 7 (29%) DUSP were observed with hyper-methylated promoters in 410 (57%) and 102 (14%) cancer cases respectively. Colorectal cancer (CRC) patients demonstrated the highest number of PTP epimutations (81%) and pancreatic cancer patients the lowest (27%) (Figure 4A). Stomach cancers presented the highest number of individuals with DUSP epimutations (23%) with lung and pancreatic malignancies the least (4%) (Figure 4B). Further analysis into the role of PTPs and DUSP revealed PTPRT is the most ubiquitously epimutated PTP, 43% of all cancer individuals showed hypermethylated PTPRT promoters. DNAJC6 (23%) and PTPRM (16%) were the second and third most epimutated PTP. PTPRT was also found to be the most epimutated PTP in 4 of the 5 cancer tissues analyzed (CRC = 78%, Stomach = 55%, Esophagus = 42% and Pancreas = 19%) and second most epimutated in Lung (21%). DNAJC6 was the most epimutated PTP in Lung (43%) (Figure 4C). CRC, stomach and esophageal cancer showed overall high levels of individuals with PTP hyper-methylated promoters. Of the top 10 most epimutated PTPs, Stomach (9/10), Esophageal (7/10) and CRC (6/10) showed >5% of individuals with epimutations in PTPs. Pancreas and Lung (2/10) presented with low epimutated PTPs (Figure4C). To a lesser extent, DUSP genes were also found to be highly epimutated (Figure 4B). DUSP26 was the gene with the highest number of individuals with hyper-methylated promoters (10%), with DUSP5 (6%), DUSP23 (3%) and DUSP15 (2%) also showing epimutations (Figure. 4D). CRC individuals showed the highest number of epimutated DUSP26 (19%) followed by Stomach (14%) and esophagus (9%). Again, lung and Pancreas showed the least (4% and 1% respectively). DUSP5 was the most epimutated DUSP in Stomach (17%), Pancreas (2%) and equal to DUSP26 in Esophageal cancer (9%) (Figure 4D).

**Figure 4:**
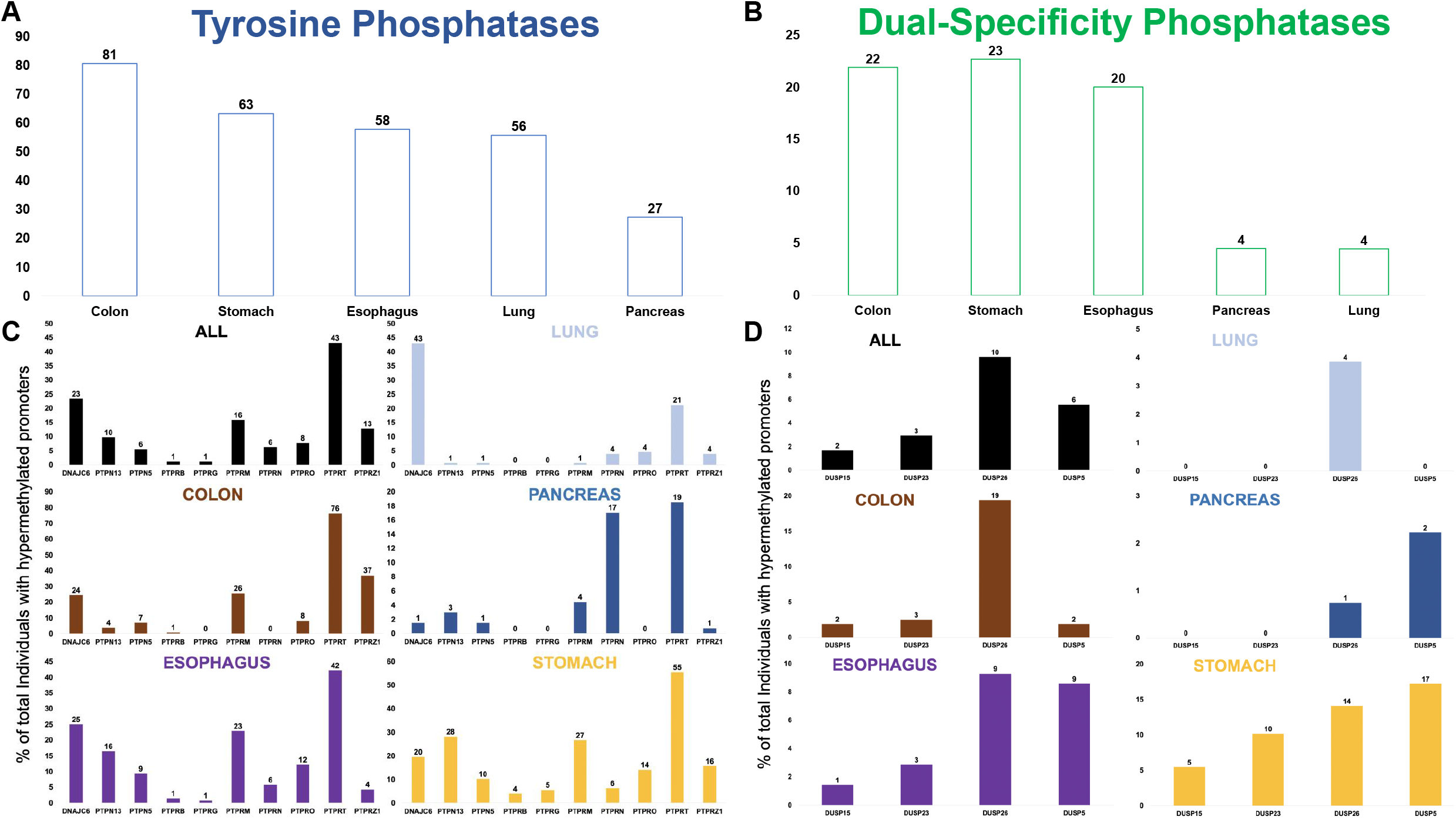
Cancer-associated epimutations are commonly observed in Protein Tyrosine (PTP) and Dual-Specificity Phosphatase (DUSP) gene promoters. A-B. Bar graph demonstrating the % of individuals in each cancer subtype that carry at least one cancer-specific epimutation in tyrosine (A, blue) and dual specificity (B, green) phosphatase gene promoters. C-D. Bar graph representing the most ubiquitous PTP (C) and DUSP (D) gene promoters that harbor cancer-specific hyper-methylated promoters. Numbers above each bar represents % of cancer individuals with epimutations in the genes highlighted on the x-axis. The genes represented in (C) and (D) were detected in at least 5 individuals.

Next, we analyzed the effect of epimutations on PTP and DUSP gene expression. 17 PTP and 7 DUSP were detected to contain at least one cancer-associated hypermethylated promoter. An initial analysis of gene expression in normal tissues was conducted using the GTEx portal (gtex.org, (Lonsdale et al., 2013)) in the five tissues analyzed in this study to determine the potential effect of cancer-associated epimutations on PTP and DUSP transcription. 9 of 17 PTPs and 5 of 7 DUSP were observed to be expressed at high levels in at least one healthy tissue type (>5 TPM) (Supplementary Figure 4A and B). Further investigation revealed that 4 PTP (PTPRM, PTPN13, PTPRG and PTPRB) and 3 DUSP (DUSP23, DUSP5 and DUSP2) were ubiquitous epigenetic outliers in all cancer cell models, highly expressed in at least one normal tissue (>5 TPM) and maintained expression in their malignant counterpart prior to segregation based on promoter DNA methylation (Supplementary Figure 5). Expression profiles from primary tumors (TCGA) and cancer cell lines (CCLE) for the 4 PTP and 3 DUSP are presented in Figure 5. In each boxplot, gene expression is partitioned by individuals with hypermethylated promoters (>0.33 average promoter beta value in TCGA and > 0.66 in CCLE vs healthy controls) (see methods for details). A significant negative correlation between promoter methylation and gene expression was observed in all genes and cell models. We expect these data to also be representative of cancer organoid cell models (Joshi et al., 2020).

**Figure 5:**
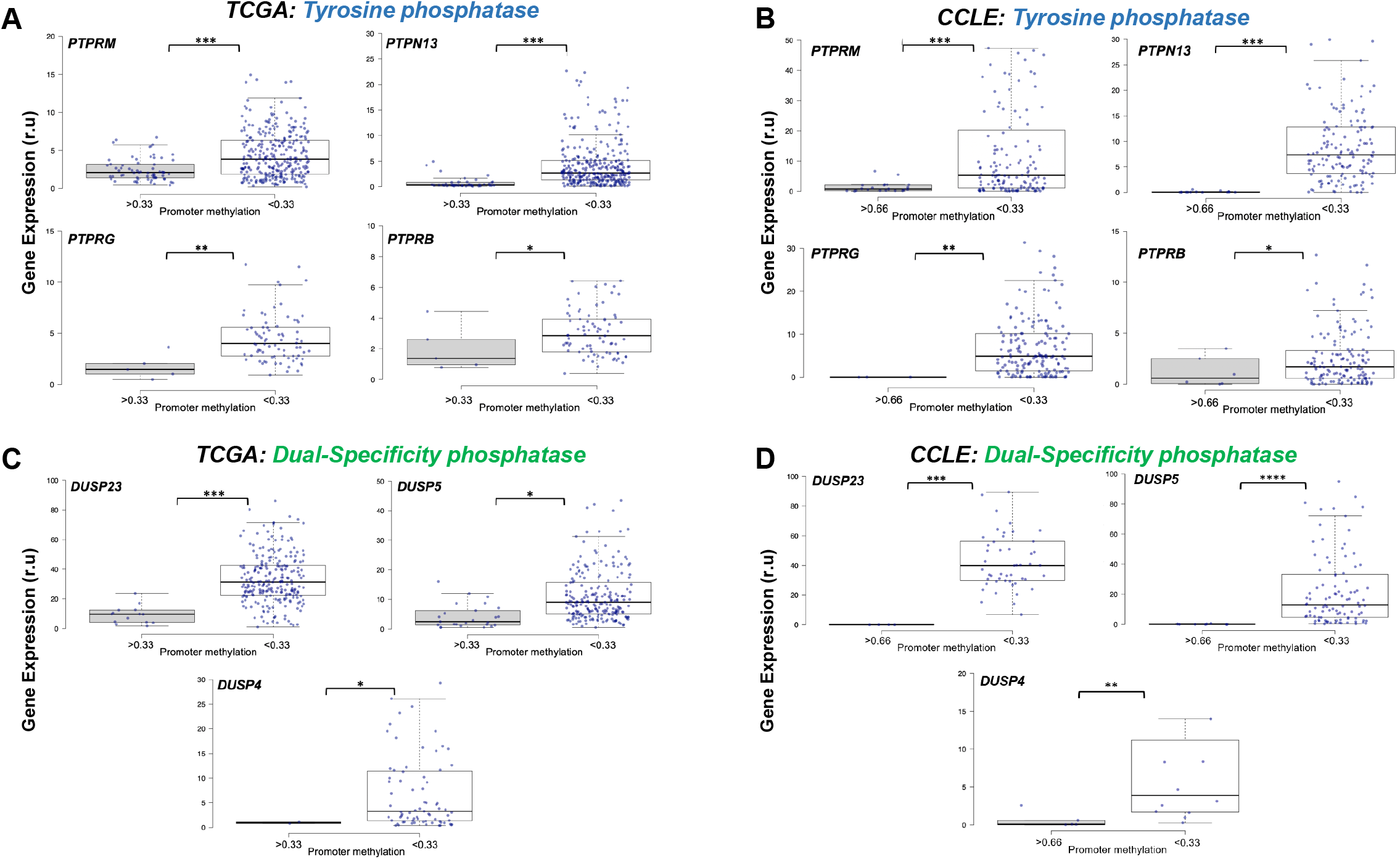
Cancer-specific promoter DNA hypermethylation in Protein Tyrosine (PTP) and Dual-Specificity Phosphatase (DUSP) are associated with gene expression silencing. A-B. Boxplots representing the effect of hypermethylated promoters of PTP genes in primary tumors (A) and cancer cell lines (B). C-D. Boxplots representing the effect of epimutations discerned in promoters of DUSP genes from primary tumors (C) and cancer cell lines (C). Primary tumor and cancer cell line gene expression data was downloaded from The Cancer Genome Atlas (TCGA) and the Cancer Cell Line Encyclopedia (CCLE) projects. Gene expression values are presented as relative units (r.u) and are specific to each project (TCGA = FPKM and CCLE = TPM). Gene names shown at the top left hand corner of each boxplot and blue dots presents individual (and cancer cell lines) expression values where DNA methylation profiles were also available. *P < 0.05; **P < 0.01; ***P < 0.001.

Although hypermethylated promoters negatively correlate with gene expression in our cancer cohort (Figure 5), one exception was observed in the gene *DNAJC6*, where an increase in promoter methylation was positively correlated with gene expression (Supplementary Figure 6A and C). DNAJC6 is the second most epimutated gene in all cancer samples and the most epimutated gene in the lung cancer cohort (43%) (Figure 2 and 4). Although a high number of epimutations were identified in our analysis cohort, several showed very low expression in their pertinent tissues. As mentioned above, PTPRT is the most epimutated gene found in this study however its expression is extremely low (or non-existent) in the 5 tissues analyzed (Supplementary Figure 6B and D) with the highest expression observed in brain tissue (Supplementary Figure 4A). This interesting finding demonstrates that epigenetic dysregulation in cancer cells occurs independent of an active transcriptional program and may provide important genomic information for other tissues.

**Figure 6:**
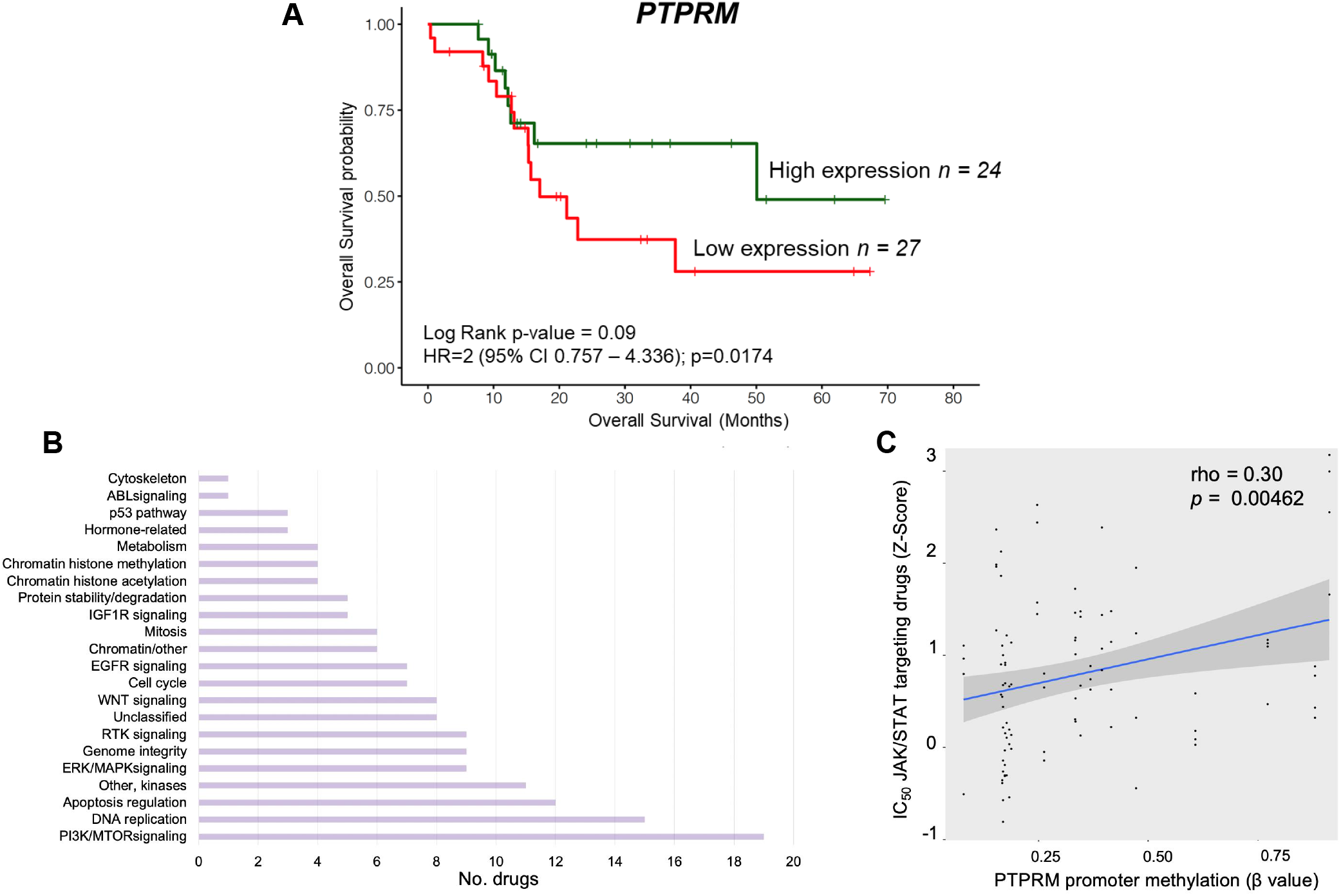
*PTPRM* reduced expression via epigenetic silencing is associated with poor survival and reduced sensitivity to JAK/STAT targeted anti-cancer therapy in pancreatic cancer patients. A. Kaplan–Meier curve showing reduced expression of PTPRM in patients with pancreatic cancer was significantly associated with shorter overall survival (HR=2; 95% CI = 0.757-4.336; P = 0.0174). Green line represents patients in the top 25-percentile of PTPRM expression and the red line, bottom 25-percentile *PTPRM* expression in the pancreatic primary tumor cohort. B. Horizontal bar graph demonstrates the cellular pathways targeted by anti-cancer drugs from GDSC2 (Genomics of Drug Sensitivity in Cancer) project. Drugs targeting “other” category were excluded. C. Scatter plot illustrating IC50 concentration of JAK/STAT pathway targeting compounds is significantly proportional to DNA methylation of the PTPRM gene promoter in pancreatic cancer cell lines (rho = 0.3, P = 0.00462).

### PTPRM epigenetic transcriptional silencing correlates with poor clinical outcome and reduced anti-cancer drug sensitivity

The role of PTP as tumor suppressors has been described in great detail in a number of tissues (Stebbing et al., 2014) and are targets for anti-cancer therapy (Köhn, 2020). In our cancer cohort, PTPRM represented the PTP with highest number of individuals with epimutations and high median expression values in healthy subjects (>5 TPM). Having demonstrated the presence of *PTPRM* promoter hyper-methylation associated transcriptional silencing (Figure 5A and B), we studied if PTPRM epigenetic loss in cancer patients had any impact on the clinical outcome in these patients. For this, we leveraged complete clinical and transcriptomic data from all individuals used in this study available in TCGA data repository. We identified that *PTPRM* gene silencing (top 25% quartile expression vs bottom 25%) in pancreatic cancer patients showed a significant association with poor overall survival probability (hazard ratio [HR] = 2; 95% confidence interval [CI] = 0.757 – 4.336; *P* = 0.0174).

PTPRM is an important component of STAT3 regulation with downstream effects on proliferation and metastasis in lung cancer (Im et al., 2020) therefore we speculated whether PTPRM epigenetically deficient cancer cells could be exploited for therapeutic purposes, specifically drugs that target the JAK/STAT cell signalling pathway in other tissues. We downloaded IC50 concentration Z scores from GDSC2 (Genomics of Drug Sensitivity in Cancer, dataset2) database (Iorio et al., 2016) for antitumor drugs that target key cancer-related cellular pathways (Figure 6B). We identified 4 compounds, AZ960 (Yang et al., 2010), JAK_8517, JAK1_8709 (Mongre et al., 2021) and Ruxolitinib (Han et al., 2018), that specifically target proteins in the JAK/STAT pathway and compared their drug sensitivity to *PTPRM* promoter methylation status in pancreatic cancer cell lines. We observed that drug sensitivity was significantly proportional to DNA methylation levels of the *PTPRM* gene promoter in pancreatic cancer cell lines (rho = 0.3, *P* = 0.00462) (Figure 6C) suggesting that *PTPRM* promoter methylation profiles maybe used as a potential biomarker for clinical treatment response in pancreatic cancer patient therapy.

## Discussion

In this study, we systematically surveyed hypermethylation profiles of Protein Phosphatase Enzymes and interacting protein (PPEIP) gene promoters to highlight the role of epigenetic marks that alter transcription of tumor suppressor genes and its effects within a clinical setting. In this regard, we detected 5007 hypermethylation events in 160 PPEIP gene promoters that were absent in healthy controls (epimutations). These epimutations were detected in 539 cancer patients (84%) across 5 tissues (colorectal, esophageal, lung, pancreatic and stomach) and may disrupt the delicate balance of protein phosphorylation in signaling networks and promote malignancies. This dysregulation of key genes is not uniform across all tissue types. By assaying epimutations in multiple tissues, we observed that stomach cancer patients accumulate the highest number of epimutations (Supplementary Figure S1 and 2) while lung cancer cases produce the least. This is an interesting finding as epimutations in PPEIP have been described in both tissues, (examples include; (Omerovic et al., 2010; Wu et al.)) although a cumulative comparison across tissues and individuals has not been previously reported. This disparity of epimutations may relate to their function defined by regulatory signatures underlying each tissue (such as active chromatin marks H3K4me3 and H3K4me1) that accompany differences in DNA methylation patterns by tissue-specific mechanisms in malignant cells (Zhang et al., 2013). Our data suggests lung cancer tumors are less susceptible to PPEIP hypermethylation inactivation that affect highly mutable signaling pathways such as PI3K-AKT, JAK/STAT and MAPK (Figure 3), and oncogenic programs that establish a favorable molecular environment for tumor suppressor inactivation likely rely on other genomic mechanisms (Greenman et al., 2007). These findings may have important clinical implications for designing treatment strategies that target PPEIP genes and their disrupted pathways.

The role of phosphatases, in particular Protein Tyrosine Phosphatases or PTP (and to a less extent, Dual Specific Phosphatases, DUSP), as tumor suppressors have been studied for decades and are emerging targets for novel technologies for oncogenic therapy (Gao et al., 2021; Stanford and Bottini, 2017). 57% of all cancer patients revealed epimutations in PTP genes and were overwhelmingly enriched in Colorectal Cancer probands (81%) as compared to pancreatic cancer (23%). Interestingly, epimutations in *PTPRT* were discerned in the most individuals despite very low expression in all tissues analyzed (Supplementary Figure 4A and 6B an D). This finding provides evidence that abnormal promoter hypermethylation mechanisms target certain tumor suppressors independent of their transcription activity. PTPRT is a negative regulator of signal transducer and activator of transcription 3 (STAT3) and therefore promoter hypermethylation events in addition to deleterious genetic mutations may be utilized as biomarkers to inform on potential neoplasm growth in multiple tissues, responsiveness to STAT3 inhibitors (Peyser et al., 2016) and predictors of standard treatments against cancer (Hsu et al., 2018). In contrast, we discerned 4 PTP and 3 DUSP genes where epimutations consistently resulted in transcription silencing in all cell models (Figure 5). All 7 genes have been implicated in several cancer-specific cellular programs as tumor suppressors and DNA methylation profiles associated with these genes may have compelling clinical implications (Stebbing et al., 2014). For example, *PTPRM* epimutations were observed in the most cancer cases of the 7 gene (16%) in all 5 tissues. PTPRM is a receptor PTP and its intracellular compartment is responsible for phosphatase activity, whereas the extracellular section serves in cell–cell and cell–matrix contact (Tonks, 2006). STAT3 inactivation is catalyzed by PTPRM dephosphorylation and leads to cancer cell death, therefore errors incurred in STAT3 dephosphorylation, such as promoter hypermethylation induced *PTPRM* transcriptional silencing, may lead to cancer initiation and progression with poor clinical prognosis (Im et al., 2020; Yu et al., 2009). Furthermore, our data demonstrated that low PTPRM expression is associated with poor overall survival (Figure 6A) and PTPRM methylation marks maybe used as biomarkers for JAK/STAT targeting anti-cancer drugs response. Additionally, the other epimutated genes showed equally intriguing roles in cancer. For example, in breast cancer, PTPN13 was reported to inhibit cancer aggressiveness by Src dephosphorylation (Glondu-Lassis et al., 2010) which is upregulated in tamoxifen-resistant ER-positive breast cancer patients (Planas-Silva et al., 2006), or DUSP4 repressed expression was identified as a mechanism of neoadjuvant drug chemoresistance and frequently depleted in chemotherapy refractory breast tumors (Balko et al., 2012). Together, this highlights the role of epimutations in PTP and DUSP genes as putative cancer biomarkers for diagnosis, prognosis and treatment response.

Interestingly, we identified one example where hypermethylated promoters resulted in an increase of transcription (Supplementary Figure 6A and C). DNAJC6 was the second most epimutated PTP gene in cancer patients and the most in lung cancer patients (Figure 4C). A literature search revealed DNAJC6 possesses oncogenic properties and promotes hepatocellular carcinoma (HCC) progression through induction of epithelial–mesenchymal transition. Overexpression of DNAJC6, as shown with increased promoter methylation in cancer subjects, was observed to enhance cell proliferation and invasion suggesting *DNAJC6* hypermethyation may be assayed as a putative biomarker for poor outcome in HCC (Yang et al., 2014).

Overall, our data illustrates the clinical relevance of DNA hypermethylation in a subset of PPEIP. We have broadened our knowledge of PPEIP epimutations across previously undescribed genes and tissues as well as providing an insight into epimutation susceptibility and distribution and its subsequent role in tumorigenesis and treatment. Considering the clinical implications, this type of phosphatase-wide analysis of promoter hypermethylation would benefit patients with other cancer types with high levels of phosphatase activity such as brain and neuroblastomas (Supplementary Figure S4). With advances in single cell sequencing technologies, targeting epimutations in phosphatase enzymes will reveal precise signaling pathways and transcriptional programs in clonal sub-populations of tumor cells previously undetectable with bulk technologies and vastly improve cancer therapeutics. Together, it’s seductive to posit that the emergence of epi-drugs that rehabilitate genes inactivated through epigenetic mechanisms (Berdasco and Esteller, 2019) hold promise for re-activating tumor-suppressing cellular programs through genes such as PTPRM, that leads to improved treatment strategies and reduction in mortality of high-risk patients.

## Materials and Methods

### Genome-wide DNA methylation array samples

A total of 729 genome-wide DNA methylation array datasets were downloaded from publicly available databases and processed in this study. Raw intensity (idats) files produced using the Infinium^®^ HumanMethylation450 (450K) BeadChip (Illumina) allowed for interrogation of 450,000 CpG’s. Raw idat files from random 500 primary tumors from The Cancer Genome Atlas (TCGA) legacy database were downloaded representing 100 samples for five TCGA projects; TCGA-COAD and READ (colorectal cancer), TCGA-ESCA, (Esophageal cancer), TCAG-LUAD (Lung cancer), TCGA-PAAD (Pancreatic cancer), and TCGA-STAD (Stomach cancer). 50 healthy control samples (10/tissue type) from each project were also analyzed in this study. Of the 50 control sets, 40 were downloaded from TCGA projects (TCGA-COAD and READ, TCGA-ESCA, TCAG-LUAD and TCGA-PAAD). We also incorporated 450K DNA methylation data from healthy stomach tissue available in GEO under accession number GSE127857. Additionally, as a test sample set to evaluate our outlier DNA methylation analysis bioinformatic pipeline in non-cancer subsets, a second, independent set of 47 healthy controls were downloaded from TCGA (TCGA-COAD (*n* = 16), TCGA-ESCA (*n* = 8), TCAG-LUAD (*n* = 16), TCGA-PAAD (*n* = 5), and TCGA-STAD (*n* = 2)) and analyzed identically to the cancer samples. 204 cancer cell line DNA was acquired from the Catalogue Of Somatic Mutations In Cancer (COSMIC) database from the Wellcome Trust Sanger Institute as detailed in (Iorio et al., 2016). Raw intensity files were downloaded corresponding to the five cancer subtypes described above; Colorectal (*n* = 49), Esophageal (*n* = 35), Lung (*n* = 61), Pancreas (*n* = 31) and stomach (*n* = 28) cancer cell lines. Finally, idat files for 25 embedded 3D cultures (organoid) representing 5 cancer subtypes were downloaded from the GEO website under the accession number GSE144213. The clinical tumor diagnoses that characterized the 5 organoid cancer subtypes were Colorectal (*n* = 11), Pancreatic (*n* = 7), Esophageal (*n* = 4), Stomach (*n* = 2) and Lung (*n* = 1) cancer.

### DNA methylation quality control, normalization and filtering

Raw signal intensity values were initially QC’d and pre-processed from subsequent idat files in R statistical environment (v3.6.1) (r-project.org) using minfi Bioconductor package (v1.32.0) (Aryee et al., 2014; Fortin et al., 2016) and processed in batches by tissue, cancer cell model and healthy samples independently. Quality control steps were applied to minimize errors and remove poor probe signals. Putative labeling errors were ascertained by examining methylation status at sex chromosomes of each individual. Duplicate samples were discerned by the SNP analysis feature in minfi. Vigorous quality control steps were performed on all samples and are detailed in (Joshi et al., 2020). Briefly, tSNE plots were produced to assess cancer subtype clustering to identify mislabeled samples. Hierarchical clustering with multi-step bootstrap resampling was performed using the R packages pvclust (Suzuki and Shimodaira, 2006) and plotted using Rtsne (LVD, 2014; van der Maaten LJP, 2008). Fifty thousand β values from across the genome in all tissue types were randomly selected in 5000 iterations to assemble and visualize inherent sample similarity. All duplicate datasets and mislabeled samples were subsequently removed from this analysis (*n* = 24). Next, problematic probes such as failed probes (detection *p* value >0.01), cross-reacting probes and probes that overlapped single nucleotide variants within +/− 1bp of CpG sites were removed. Background correction and dye-based normalization was performed using ssNoob algorithm (single-sample normal-exponential out-of-band). Lastly, chromosomes X and Y were removed from the analysis. In total, 705 cancer samples, 42 healthy controls and 47 healthy test controls were used in our final analysis. All QC, normalization and filtering steps mentioned above were performed separately for each cancer subtype and cell model. The control test samples were ran separately to the tumor samples. A breakdown of all samples can be found in Supplementary data S2. Final DNA methylation values for each CpG, represented as β-values with 1 representing fully methylated CpG and 0, fully unmethylated CpG were used for analysis. Downstream analyses were performed under R statistical environment (v3.6.1).

### Gene expression profiles

In order to assess the putative consequence of DNA methylation on the expression status of all TCGA individuals that harbored outlier DNA methylation at promoters of Protein Phosphatase Enzymes and interacting proteins (PPEIP) and controls, RNA-seq Fragments Per Kilobase of transcript per Million mapped reads (FPKM) values were downloaded using the R Bioconductor package TCGAbiolinks (Colaprico et al., 2016; Silva et al., 2016) for all primary tumors and healthy tissues. Transcription profiles were only downloaded from patients where DNA methylation data is also available. For cancer cell lines, gene expression profiles (Transcripts Per Kilobase Million or TPM) used for the DNA methylation analysis were downloaded directly from the Cancer Cell Line Encyclopedia (https://portals.broadinstitute.org/ccle) website. No analysis was performed between the two different types of datasets (eg FPKM vs TPM); relative expression profiles were only assessed within their specific projects, analysis pipelines and expression units.

### Clinical data

Extensive clinical information for all samples from the TCGA project (primary tumors) were retrieved from TCGA using TCGAbiolinks and integrated into R statistical environment (v3.6.1) for overall survival analyses.

### Data analysis pipeline

A curated list of 726 unique Protein Phosphatase Enzymes and interacting proteins (PPEIP) genes was downloaded from Ensembl (http://www.ensembl.org). PPEIP genes were defined as those genes that encode a protein that has been with previously described phosphatase activity and/or encode proteins that form PPEIP subunits, interact with PPEs that potentially alter their function. 523 genes had at least one 450K probe associated to its promoter and subsequently used for analysis. The full list can be found in Supplementary data S1. Promoter associated probes were defined as 450K probes that measure CpG DNA methylation levels within 1500bp and 200bp from the transcription start site as well as the 5’UTR and 1^st^ exon of all PPEIP genes. Probe annotation (such as those located in promoters, CpG island/shores, enhancers, etc) is provided by Illumina’s 450K manifest file. Average promoter methylation per gene was calculated in 705 cancer samples (including primary tumors, cancers cell lines and organoids), 42 baseline healthy control subjects and 47 test controls. A preliminary screen of PPEIP genes that demonstrated >0.33 average promoter beta values in healthy tissues were discarded. Epimutations or outlier individuals that harbored hypermethylation marks in PPEIP promoters were defined as >0.66 average beta values in cancer cell lines and >0.33 average beta values in primary tumors and organoid samples as compared to baseline healthy controls. Each cancer tissue type was only compared to its corresponding healthy tissue counterpart average promoter methylation value. This analysis and all downstream applications were performed using R statistical environment. A final list of average promoter methylation values for all 523 PPEIP genes in all 705 cancer cases and 42 baseline healthy controls can be found in Supplementary data S7.

### Epimutation tissue distribution and statistical analysis

In an even epimutation distribution (*n* = 5036), one would expect the number of epimutations to be proportional to the number of individuals per tissue set. To test this, we calculated the observed epimutation distribution ratio (OEDR) for each tissue. The equation is shown below. The sample share is the fraction of individuals in each tissue tested for epimutations. Epimutation share per tissue is the fraction of epimutations observed in each tissue and expected to be the same value as the tissue sample share in an even distribution model. The observed epimutation distribution ratio (OEDR) is the tissue sample share / epimutation share. For visualization purposes, the OEDR was converted to log2 values. This was calculated for each tissue.

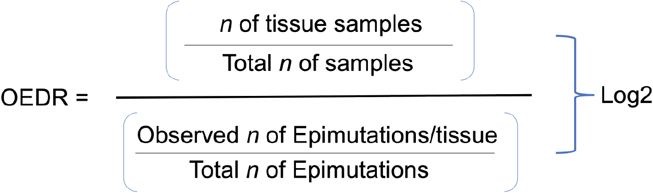

For Parametric tests, Pearson correlation and unpaired students t-tests were applied and for non-parametric data distributions, Spearmans’ rank correlation and Wilcoxon rank test were calculated. For enrichment analyses, Fishers exact and Chi-squared tests were used. Data normality was assessed using a Shapiro-Wilk test. Linear regression models were used to estimate statistical relationships between DNA methylation and drug sensitivity. Kaplan-Meier plots and Log-rank (Mantel-Cox) tests were used to estimate overall survival (OS) in pancreatic cancer individuals with outlier *PTPRM* expression. Individuals in the top 25% quartiles of *PTPRM* expression were considered as “high expression” and low expression the bottom 25% quartile. The extreme 25% quartiles were used as standard cut off points and allow for robust statistical analysis performed through Cox proportional hazards regression models. All statistical analyses were carried out with the R statistical environment (v3.6.1) and p values < 0.05 were considered as statistically significant.

### Pathway and transcription factor binding analysis

Biocarta, KEGG and curated wiki pathway analyses were carried out to highlight cellular features and cancer networks that are putatively affected by hyper-methylated PPEIP gene promoters. Transcription factor (TF) gene target examination using Encode data was also performed to inform on TF regulatory networks. All analyses were performed using the R package Enrichr (Kuleshov et al., 2016).

### Drug sensitivity

IC_50_ Z scores corresponding to drug sensitivity were downloaded from the Genomics of Drug Sensitivity in Cancer database(https://www.cancerrxgene.org/). This repository provides drug response data and genomic markers of sensitivity for 809 cancer cell lines and 198 compounds as part of their GDSC2 data release (Iorio et al., 2016). All cancer cell lines pertinent to this study were downloaded and processed according to tissue type.

---

**Supplementary figure 1:**
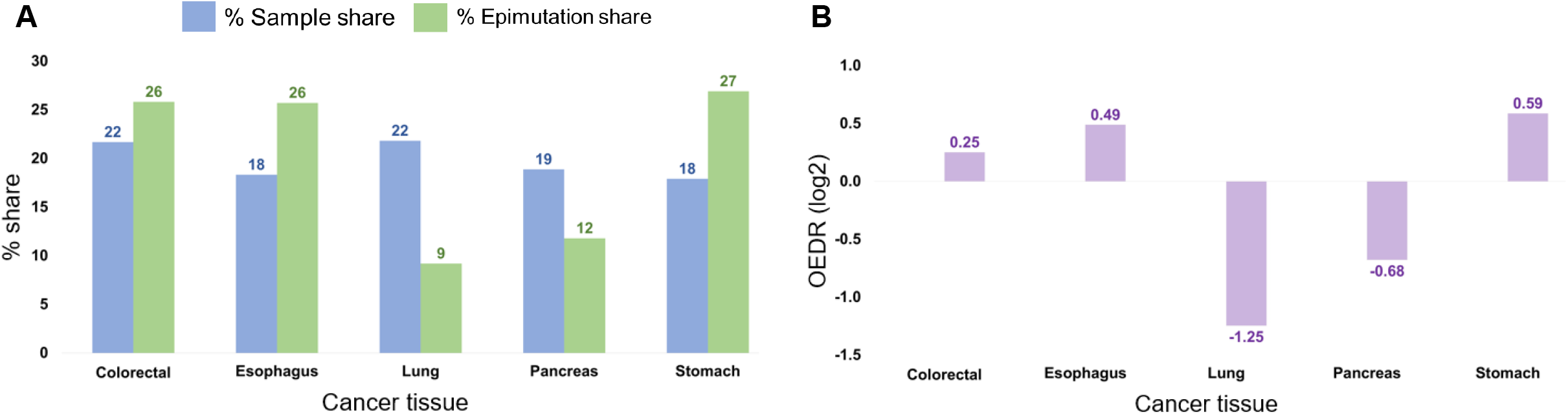
Analysis of the distribution of PPEIP epimutations across cancer sample types. A) Bar graph to show the percentage of PPEIP epimutations detected in each tissue vs total sample distribution. B) Observed Epimutation Distribution Ratio (OEDR) demonstrates the number of epimutation detected vs the expected value (0). Data reveals a disproportionate distribution of epimutations across tissue types.

**Supplementary figure 2:**
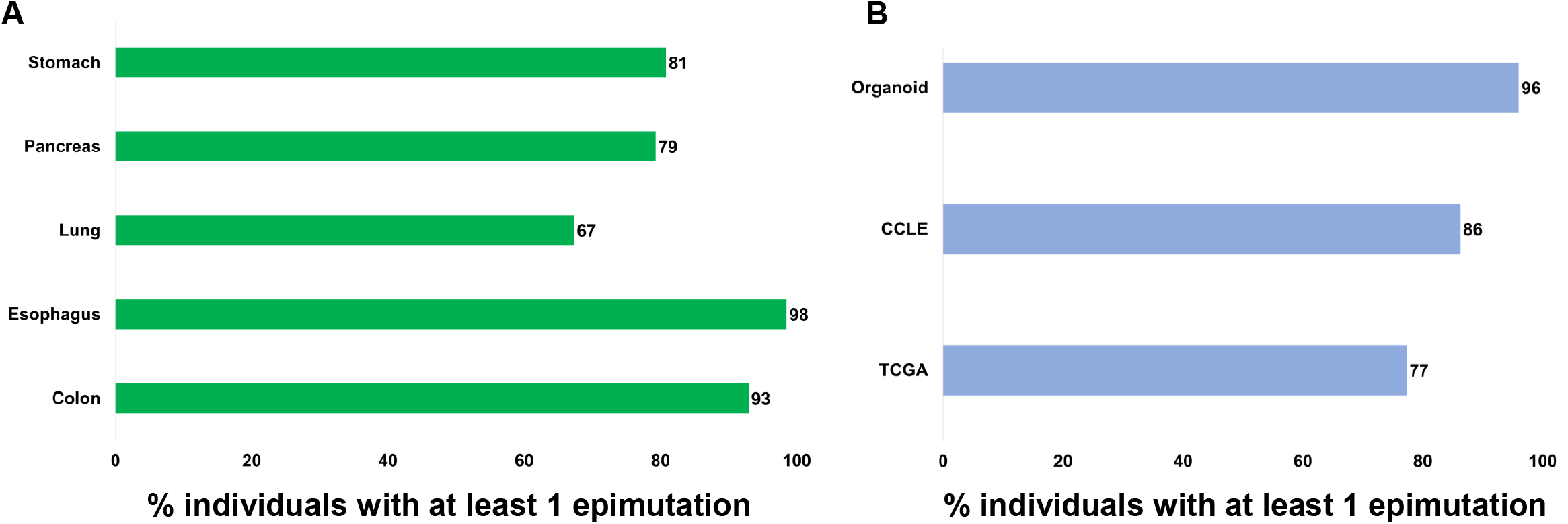
Breakdown of epimutation percentage detected in (A) cancer tissue types and (B) cancer cell models. Organoids represents 3D embedded cancer cell models, CCLE, cancer cell lines from the Cancer Cell Line Encyclopedia, TCGA, The Cancer Genome Atlas.

**Supplementary figure 3:**
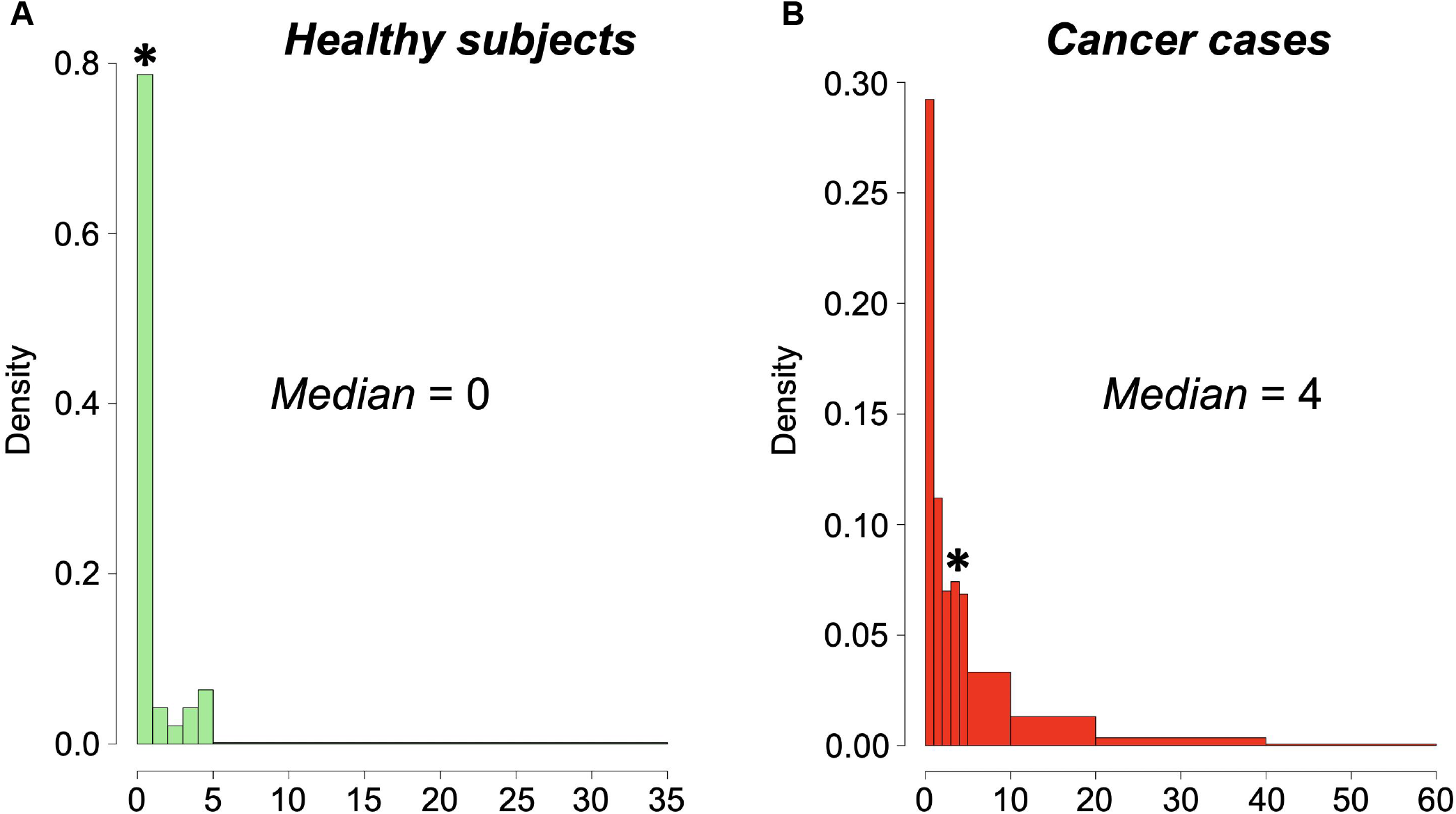
Histogram showing the number of hypermethylated PPEIP promoters identified in healthy control subjects vs Cancer cases. A) The number of PPEIP epimutations in healthy individuals in the population control cohort vs control baseline samples. B) Epimutations in each individual tumor sample. Note * represents the median value of epimutations per cohort.

**Supplementary figure 4:**
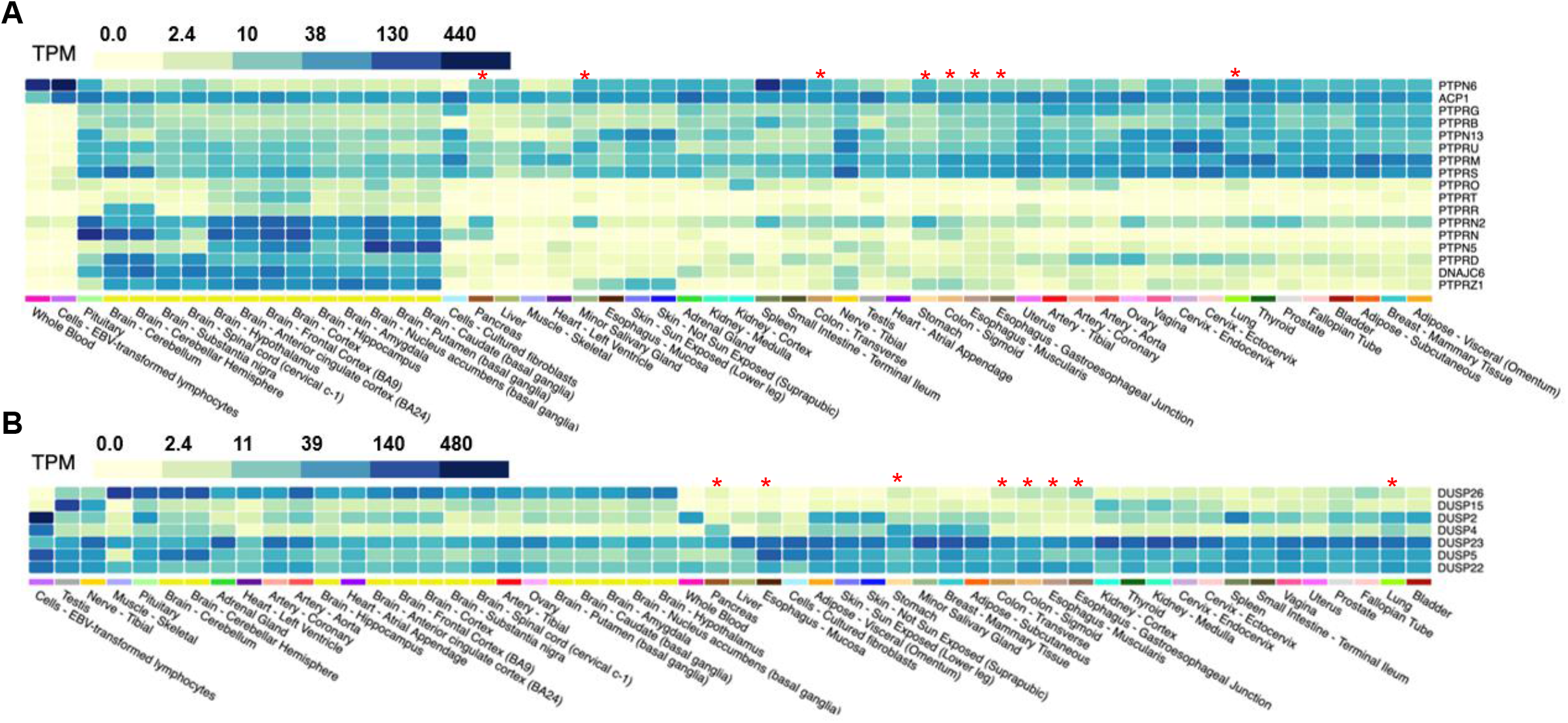
Color coded visualization of PTP and DUSP expression values in healthy tissues. A) 17 PTP and B) 7 DUSP genes were discerned with epimutations in this study and their expression values in a non-cancer population from the GTEx project presented here. All values are represented as Transcripts per million (TPM). * denotes the five tissues and subtypes analyzed in this study.

**Supplementary figure 5:**
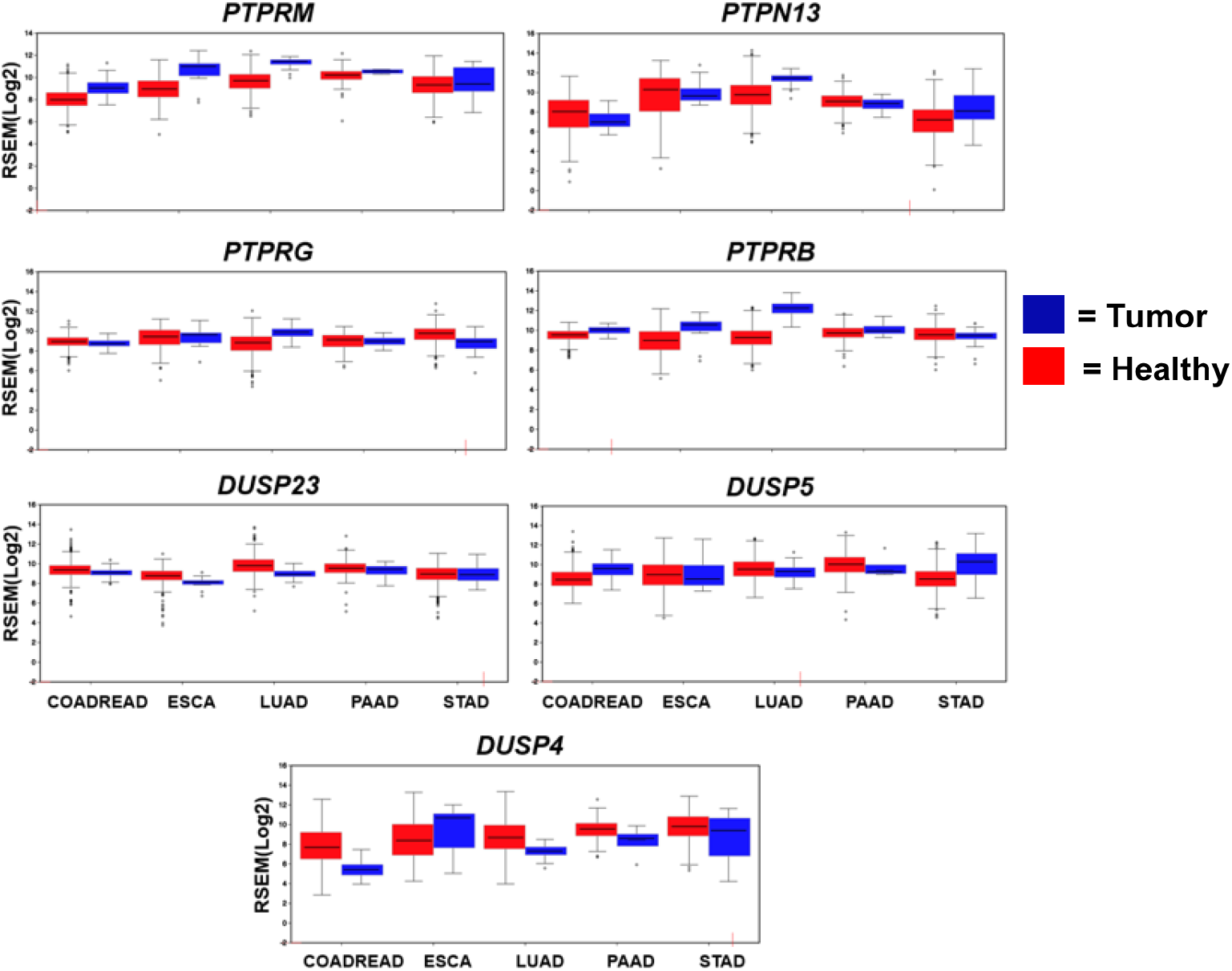
Expression of ubiquitously epimutated PTP and DUSP genes comparing normal vs cancer tissues. Transcript profiles from five TCGA projects utilized in this study (COADREAD = Colon/Rectum adenocarcinoma, ESCA = Esophageal carcinoma, LUAD = Lung adenocarcinoma, PAAD = Pancreatic adenocarcinoma, and STAD = Stomach adenocarcinoma) shows the expression of highly epimutated PTP AND DUSP genes detected in multiple cell models independent of promoter methylation status. Transcript abundance shown as RSEM (RNA-Seq by Expectation Maximization) log2 values.

**Supplementary figure 6:**
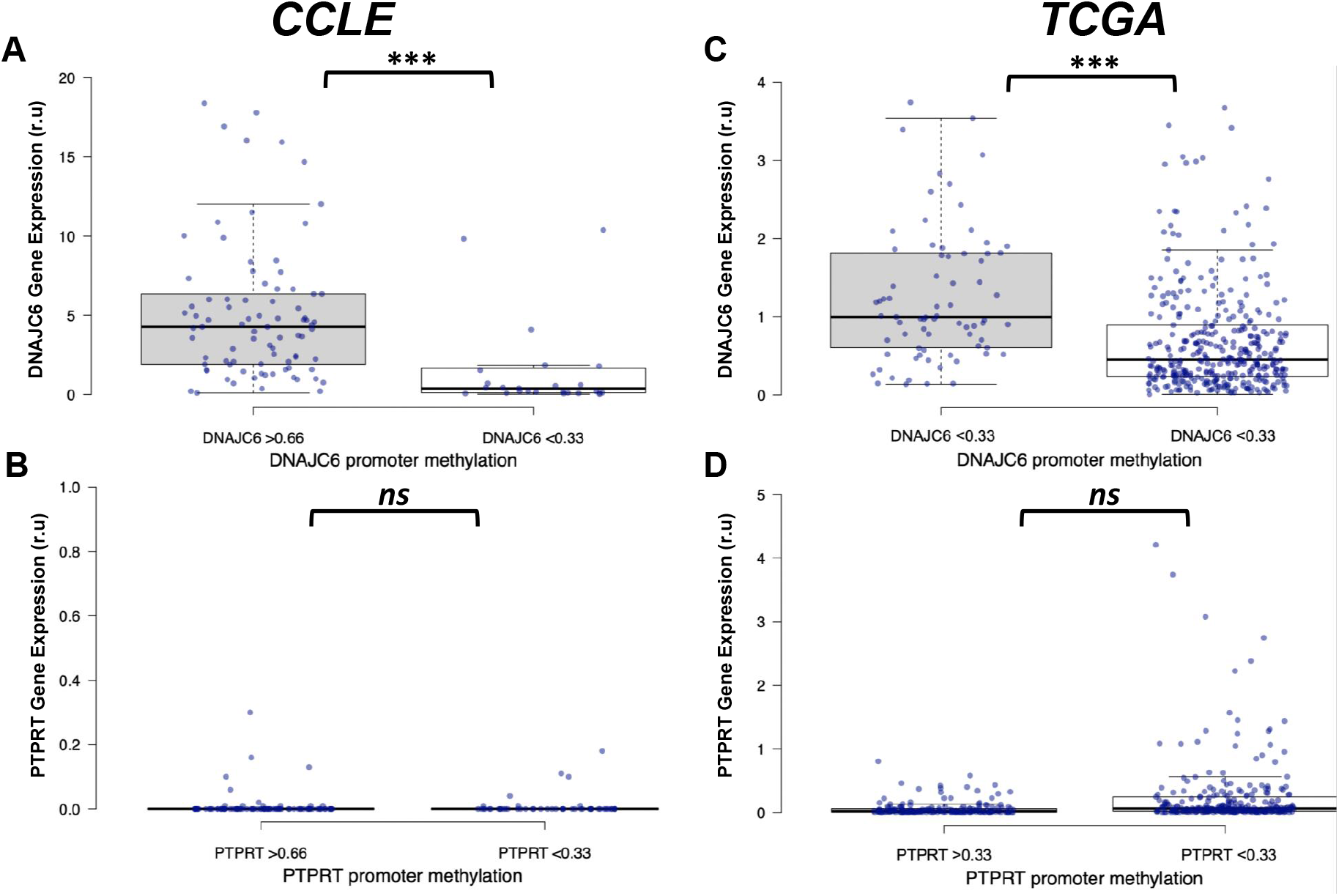
Gene expression comparison of *PTPRT* and *DNAJC6* in individuals with hypermethylated promoters vs healthy controls. A and B) represent expression profiles in cancer cell lines. In the CCLE cohort, hypermethylated promoters were those with average >0.66 beta values vs healthy baseline controls (<0.33). C and D) demonstrate expression profiles in primary tumors. Hypermethylated promoters in the TCGA subset were defined as average >0.33 beta value vs baseline beta values. Each purple dot represents expression values provided by CCLE (TPM) and TCGA (FPKM) for each individual. *** *P* < 0.001, n*s* = not significant.

## Notes

### Competing Interest Statement

ME is a consultant of Ferrer International and
Quimatryx. The authors declare that they have no conflict of interest.

### Summary of Updates

Further clarity on acronyms

